# A modular pipeline for evidence-integrated genome annotation across species: a case study on *Schmidtea mediterranea*

**DOI:** 10.1101/2025.04.16.649076

**Authors:** Anastasiia Zaremba, Małgorzata Marszałek-Zeńczak, Annasha Dutta, Anna Samelak-Czajka, Paulina Jackowiak

## Abstract

Despite advancements in genome annotation tools, challenges persist for non-classical model organisms with limited genomic resources, such as *Schmidtea mediterranea*. To address these challenges, we developed a flexible and scalable genome annotation pipeline that integrates short-read (Illumina) and long-read (PacBio) sequencing technologies. The pipeline combines reference-based and *de novo* assembly methods, effectively handling genomic variability and alternative splicing events. To improve splice site detection accuracy, DeepSplice deep learning predictions are used. Functional annotation is conducted to filter out low-confidence transcripts and ensure biological relevance. Applying this pipeline to the asexual strain of *S. mediterranea* revealed thousands of previously undescribed putative genes and transcripts, and improved the existing gene models, highlighting its utility in annotating complex, underexplored genomes. The modularity and comprehensiveness of our pipeline ensure its adaptability for genome annotation across diverse species, making it a valuable tool for annotating genomes of non-model organisms and supporting broader genomic research. The source code and implementation details are available at https://github.com/Norreanea/SmedAnno.

## Background

Understanding the genetic makeup of organisms is crucial for unraveling the complexities of their biology, making genome annotation a foundational aspect of molecular and genomic research. By precisely identifying coding regions, non-coding elements, and regulatory sequences, genome annotation transforms raw genomic data into a practical blueprint that opens up possibilities for studying living organisms at the molecular level.

Existing genome annotation methods can be broadly categorized into *ab initio* [1], homology-based [2], evidence-based (reference-based) [3], and hybrid approaches [4–6], each leveraging different strategies and tools to predict and validate genomic features. Each category has its unique advantages and challenges. *Ab initio* methods, which predict genes solely from intrinsic sequence signals without external evidence, are capable of identifying novel genes; however, they tend to produce a higher rate of false positives and inaccurate gene models—such as overpredicted exons, missed genes, and incorrect exon–intron boundaries—especially when applied to complex or poorly characterized genomes. Homology based methods use evolutionary conservation to achieve high accuracy for well conserved genes but often miss novel or rapidly evolving genes that lack close homologs. Evidence based approaches utilize transcriptomic or proteomic data to support gene models, thereby enhancing annotation quality for expressed genes, yet they are inherently limited to regions where experimental data exist and are as robust as the underlying evidence. Hybrid approaches combine intrinsic signals with external evidence in an effort to balance sensitivity and specificity.

To streamline the annotation process, several genome annotation platforms and pipelines have been developed. For instance, the RefSeq eukaryotic annotation pipeline at NCBI [7] employs highly curated alignment-based evidence and gene prediction models, while another recently reported pipeline TAGADA [8] aims to integrate evidence from multiple data sources for eukaryotic genome annotation. Web-based platforms such as Galaxy Genome Annotation (GGA) [9] provide an accessible interface for constructing annotation workflows, and tools like Prokka [10] offer rapid prokaryotic annotation that automates the integration of gene prediction and functional annotation, generating standardized output formats compatible with GenBank. Additionally, PAGIT (Post Assembly Genome Improvement Toolkit) [11], the community-driven Nextflow-based approach nf-core GenomeAnnotator [12], and other pipelines [13–15] expand the available toolkit.

Despite these advancements, applying genome annotation methods to non-model or non-classical model organisms presents significant challenges. Due to limited reference data, tools must rely more heavily on *ab initio* methods, which—as noted above— are prone to errors, particularly for complex genomes, and often lack sufficient training data. Repetitive sequences and the presence of multiple isoforms complicate the annotation process. Genetic diversity and evolutionary distance from well-studied species can impede the identification of orthologous genes and conserved pathways, while limited experimental data often constrain the ability to validate or refine gene models. Such constraints are further compounded by annotation artifacts—including fragmented and chimeric transcripts—that can obscure true biological functions and introduce errors into downstream analyses.

These challenges are particularly relevant to research on the sexual (S2F2 clonal line) and asexual (CIW4) strains of *S. mediterranea*, a highly regenerative flatworm [16,17]. Currently, reference genome sequences and comprehensive annotations are available primarily for the sexual strain, including contig-level genome assemblies such as SmedGD_c1.3 [18,19], dd_Smes_g4 [20,21], chromosome-scale genome assemblies schMedS2 [22] and recent schMedS3 [23], as well as corresponding annotations like SmedGD_c1.3, high-confidence SMESG, schMedS1, schMedS2, schMedS3_1 and schMedS3_2. Chromosome scale assemblies of the sexual strain indicate that sexual and asexual *S. mediterranea* share high genomic similarity—they are both diploid with four pairs of chromosomes. Although the asexual strain possesses a chromosomal translocation between chromosomes 1 and 3, these reference sequences and annotations offer a strong starting point for transcriptome analysis. Nonetheless, given potential strain specific transcripts and structural variations, further refinement of the asexual strain’s transcript set is essential [24]. Additionally, the asexual strain exhibits a simplified reproductive strategy [25–28] and it has been widely used in numerous studies, with extensive data deposited in public repositories. As of January 2, 2025, there are 122 DNA and 1,694 RNA datasets on NCBI [29] for this strain, the majority of which are short-read data. In comparison, for the sexual strain, there are only 18 DNA and 87 RNA datasets. The predominance of RNA-seq data highlights the extensive transcriptomic resources available for the asexual strain, which offers a strong foundation for gene expression and functional studies and underscores the potential for leveraging RNA sequencing to enhance genome annotation efforts. However, limited detailed reference resources for asexual strain remain a significant challenge. Currently, the scientific community must rely mostly on reference transcriptomic data—such as dd_Smed_v6.pcf.contigs.fasta [21] (a transcriptome assembly generated from RNA Seq data) and a contig based genome assembly and annotation SmedAsxl_genome_v1.1 [19]. Beyond that, existing transcriptome assemblies are fragmented and incomplete, lacking the high-resolution insights offered by newer sequencing technologies. To bridge this gap, our study aims to implement a sophisticated bioinformatics pipeline to improve annotated genes and transcripts in asexual *S. mediterranea* using the existing chromosome-scale genome of the sexual strain (schMedS2) as a reference and in-house generated comprehensive collection of transcriptomic data for asexual worms. Our dataset includes both short-read and long-read data, providing a significant advantage in resolving complex isoforms. Addressing these challenges requires the integration of various sequencing technologies and advanced bioinformatics tools. Short-read sequencing platforms, such as Illumina, offer high accuracy and depth of coverage, a low error rate, which is essential for identifying single-nucleotide polymorphisms, small indels, and importantly, splice sites—ensuring accurate distinction of exon-intron boundaries [30]. Conversely, long-read technologies like PacBio provide valuable information on structural variants and complex transcript isoforms, enhancing the resolution of genome assemblies [31,32]. Combining these technologies benefits from their complementary strengths, enabling a more comprehensive and accurate genome annotation [4].

Here we present a universal, modular, and adaptable pipeline for genome annotation to process and interpret the vast amounts of sequencing data generated, that integrates both short- and long-read sequencing data, employs state-of-the-art bioinformatics tools for transcriptome assembly and genome characterization, and incorporates robust methods for functional and homology annotation. By doing so, we aim to advance genome annotation efforts within *S. mediterranea* and to provide a scalable framework applicable to other non-model organisms, fostering broader genomic research and discovery.

## Methods

### Planarian culturing and sample collection

Asexual *S. mediterranea* were cultured as described in [33] and fed chicken liver twice weekly, and were used either in wild-type form or after RNAi treatment, as detailed in Table S1. Animals were collected, immediately flash frozen in liquid nitrogen to preserve RNA integrity, and stored at −80 °C until RNA isolation.

### Dataset details

Full *S. mediterranea* dataset specific details including experimental treatments, RNA extraction, quality control, are provided in Table S1 and in the associated GEO submissions. Table S1 lists transcriptomic cDNA libraries from the asexual CIW4 strain of *S. mediterranea*: 3 Illumina short-read runs (2 on NextSeq 550, 1 on NovaSeq 6000) and 3 PacBio long-read runs (1 on Sequel II, 2 on Sequel IIe). Publicly available short-read Illumina RNA-seq data for *Arabidopsis thaliana* (PRJNA477336) and *Caenorhabditis elegans* (CRA008476) were retrieved from https://www.ncbi.nlm.nih.gov/ and https://ngdc.cncb.ac.cn/gsa, respectively.

### Validation with simulated data

A synthetic genome was created using a custom Python script, which constructed a single chromosome spanning 2 mega base pairs (bp). This genome included 1,000 genes randomly distributed along the chromosome, each comprising between 2 and 20 exons with lengths ranging from 100 to 1,000 nucleotides (nt) and introns of 100 to 1,000 nt. Paired-end short-read RNA-seq data were simulated using ART Illumina tool, generating 100 bp reads at 10× coverage across six samples. Additionally, long-read sequencing data were produced using PBSIM tool with 150 bp reads at 10× coverage for one sample. To evaluate the pipeline’s ability to handle incomplete annotations, a random subset of genes was removed from the annotation file using the custom bash script, resulting in a GTF file with missing gene information.

### Reference annotation polishing

The reference genome schMedS2 [21] was utilized for this study. High-confidence annotations for this genome were obtained from two sources: SmedSxl smed_chr_ref_v1 UCLA (SIMRBase (https://simrbase.stowers.org)) and schMedS2_PlanMine_high-confidence_liftover (PlanMine [21]). Both annotations are compatible with the genome sequence; however, due to minor differences between them, the genes were first unified, and the annotation was carefully reviewed and corrected. Through detailed analysis and comparison, 27,676 SMEST transcripts and 21,332 SMESG genes were identified as the most confident and consistent across both sources. Next, 169 transcripts with identical exon structures using the AGAT tool [34] were removed to eliminate redundancy and enhance annotation quality.

### Benchmarking against BRAKER3

#### Input data

Illumina RNA-seq reads trimmed with TrimGalore were aligned to reference genome with STAR v2.7.11a [3]. For *A. thaliana* and *C. elegans*, the following reference genomes and annotations were used: GCF_000001735.4_TAIR10.1 [35,36] and PRJNA13758/ WBcel235 (WBPS19) [37,38], respectively.

#### BRAKER3 test

BRAKER3 v2.1.8 [39]was launched in “RNA-seq only” mode: braker.pl --genome genome.fa --bam star.bam --softmasking --threads 8.

#### SmedAnno test

Four runs were produced: no guide and no TransDecoder [40] (-TD) annotation added, no guide but +TD, DeepSplice [41] guidance with -TD, DeepSplice guidance with +TD. All used StringTie2 v2.1.1 and omitted long-read input.

#### Structural accuracy

Predictions were compared to references with gffcompare v0.12.6 [42]. Sensitivity, precision and F1 score were extracted at exon, intron and transcript level.

#### Completeness

BUSCO v5.7.1 [43] was run in transcriptome mode (-m tran) with the following lineages: Embryophyta_odb10 (*A. thaliana*), Nematoda_odb10 (*C. elegans*) and Metazoa_odb10 (*S. mediterranea*).

### StringTie2 parameter optimization

The transcriptome assembly was evaluated using the SIRV-set 4 RNA spike-in controls to assess its accuracy, with specific StringTie2 parameters (-c, the minimum reads per bp coverage to consider for multi-exon transcript, and -f, the minimum isoform fraction) being tested (Table S2). Five combinations of StringTie2 parameters were used: default settings (-c 1; -f 0.01), a stricter isoform fraction filter alone (-f 0.02), two higher coverage thresholds (-c 2 and -c 5), and a mixed setting (-c 1.5, -f 0.02).

### Assembly evaluation and comparative analysis

Completeness was assessed with BUSCO [43] (v5.7.1) with a Metazoa benchmark of 954 conserved genes, while assembly quality metrics were obtained using rnaQUAST [44] (v2.3.0) and Transrate [45] (v1.0.3). Benchmarking procedures were conducted by comparing the performance of the pipeline against existing annotations, with metrics such as the number of correctly identified transcripts, assembly scores, and completeness indicators being analyzed.

### ncRNA identification

All ncRNA sequences were downloaded from RNAcentral [46] and supplemented with 457 tRNA sequences from an external publication [47]. The sequences were linearized and deduplicated using seqkit rmdup. The initial dataset contained 27,390,440 sequences, which was reduced to 27,317,511 after deduplication. A BLASTn[2] search was conducted to identify homologous sequences within the *S. mediterranea* nuclear and mitochondrial genomes. The following parameters were used: -query: 27,317,511 ncRNA sequences; -db: *S. mediterranea* genome (nuclear + mtDNA), -evalue: 1e-20, -word_size: 15, -qcov_hsp_perc: 90, -dust: no. The output was then processed to extract the original sequence length of hits for further filtering. A custom Python script (Table S3) was used to classify ncRNA types based on query ID patterns. Hits were retained if alignment coverage met predefined thresholds (95% for tRNA, 90% for miRNA, 70% for lncRNA, 90% for rRNA, 80% for other ncRNA). Then another custom R script (Table S3) was used for converting to gtf format, refining and merging overlapping regions to ensure only non-redundant ncRNA annotations were retained.

## Analyses

### Selection and validation of transcriptome assembly approach

One of the key steps in genome annotation is the assembly of the transcriptome. In our efforts to enhance genome characterization for the asexual strain of *S. mediterranea*, we employed StringTie2 [4], a widely recognized tool for transcriptome assembly. StringTie2 is versatile, supporting both short-read and long-read sequencing datasets, which makes it well-suited for comprehensive transcriptome analysis. However, StringTie2 has multiple versions, each offering distinct features and performance enhancements. Therefore, before integrating StringTie2 into our pipeline, we thoroughly optimized its parameters to ensure the highest accuracy and reliability in transcript predictions. To this end, we employed two complementary strategies.

In the first strategy, we generated an *in-silico* dataset to simulate genomic sequences with corresponding full and incomplete annotations and short-read RNA-seq data (see Methods section). This synthetic data allowed us to create scenarios with known simple transcript structures, enabling a comprehensive evaluation of StringTie2 performance. Reads were aligned to the *in-silico* reference genome using STAR. We then compared two commonly available versions of StringTie2, 2.1.1 and 2.2.1, which allowed us to assess version-specific performance improvements. We evaluated the results of the assembly step, by analyzing transcript identification rates for both StringTie2 versions at base, exon, locus and transcript levels using gffcompare. Our results indicated that StringTie2 version 2.1.1 provided a higher identification rate of short-read data in reference-based mode than its descendant (Figure 1A), while there were no differences in *de novo* mode. Although the StringTie2 documentation does not explicitly state that version 2.1.1 is more sensitive for short-read data, the combination of release note details and our experimental observation suggested that less stringent filtering of version 2.1.1 results in a higher count of assembled isoforms. Meanwhile, version 2.2.1 has been optimized for hybrid-read assembly (with the --mix option) and incorporates more conservative measures to reduce false positives—which can lower the total transcript count when only short-read data are used. This trade-off between sensitivity and precision is common in transcript assemblers. When maximizing the number of detected isoforms from short reads is a priority, 2.1.1 may be advantageous, whereas 2.2.1’s improvements (including its ability to process mixed-read data) make it preferable for datasets that include long reads. It should be noted that these version choices are based on their prevalence at the time of our study, and although our benchmarking indicates reliable performance, further testing in diverse datasets may be beneficial.

**Figure 1.**
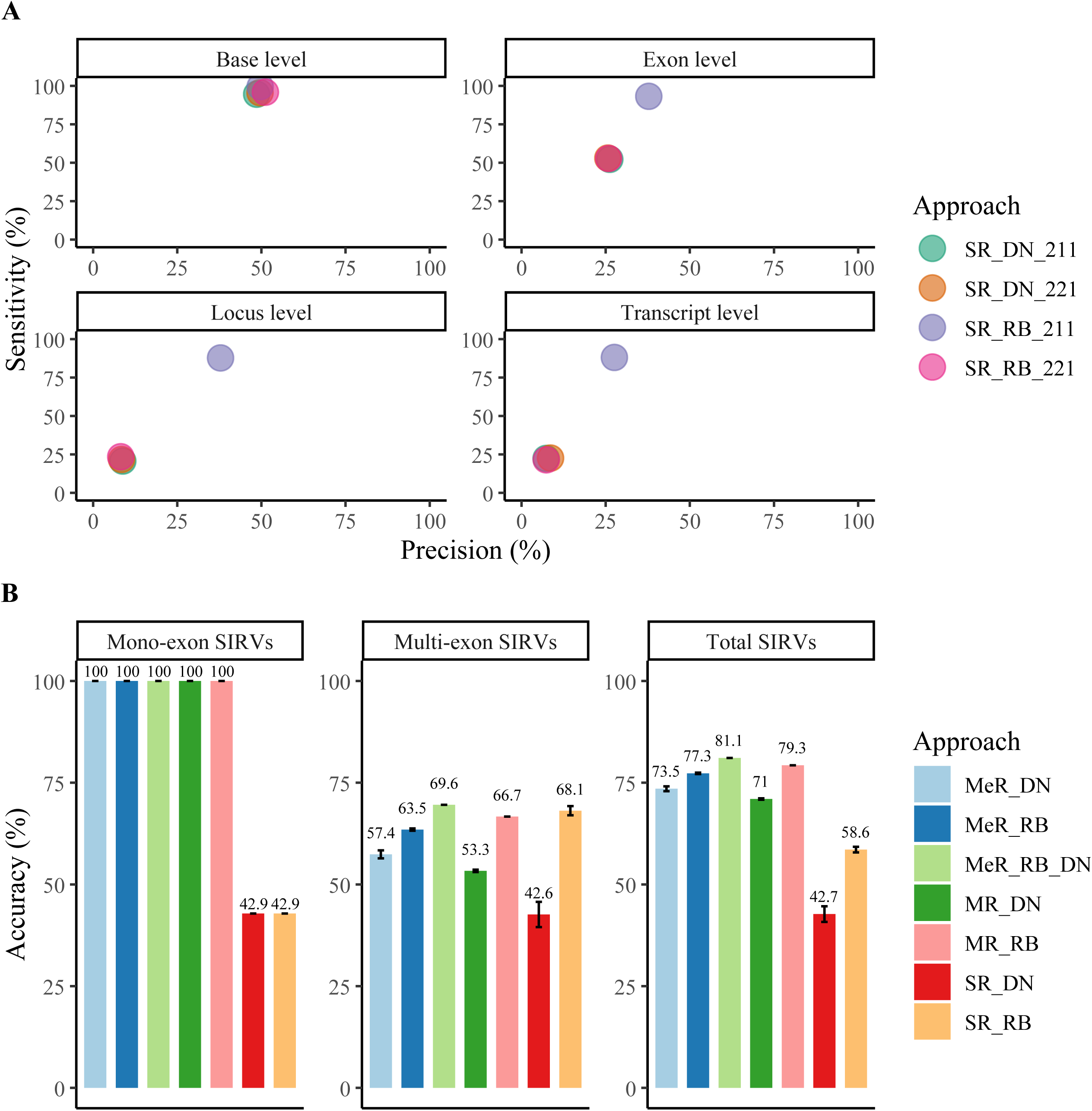
Identification rates and accuracy metrics for *in-silico* and SIRVs datasets. (A) - Transcript identification rates using *in-silico* data. Circles depict the four approaches tested for short read (SR) data, using *de novo* (DN) or reference-based (RB) assembly and StringTie2 version 2.1.1 (211) or version 2.2.1 (221). (B) - Accuracy metrics of transcript isoform identification using SIRVs on SR, mixed reads (MR) and merged reads (MeR) approaches. Whiskers on the bar plots represent the variability in accuracy across five different StringTie2 parameter configurations tested.

In the second strategy, we used Spike-In RNA Variants (SIRVs), a commercially available collection of RNA spike-in controls, which were added to the RNA isolated from planarians (see Table S1) prior to the preparation of sequencing libraries. These SIRVs included three distinct modules — ERCC Mix 1 (a set of 92 synthetic RNA molecules ranging between 250 and 2,000 nt in length to resemble natural eukaryotic transcripts), Isoform Mix E0 (referred here as multi-exon SIRVs), and long SIRVs (referred here as mono-exon SIRVs) — each designed to mimic specific aspects of transcriptome complexity, including transcript abundance, isoform diversity, and transcript length. RNA-seq reads were aligned to the reference SIRVs sequences using STAR for short reads and Minimap2 for long reads. We then employed StringTie2 using the “insufficient annotation” as the reference provided by the manufacturer of SIRVs (SIRV_I), where some SIRVs that are actually present in the mixes are not annotated, and compared the prediction to the correct annotation of all SIRVs (SIRV_C). Such a comparative analysis demonstrated a proper reconstruction of missing spike-in transcripts. Then, focusing on the mixture of synthetic RNA isoforms Iso Mix E0 category, we systematically assessed the performances of seven assembly approaches for transcript variant detection, implementing different combinations of read types—short and mixed—and assembly modes—reference-based and *de novo*. These approaches included short reads in reference-based mode (SR_RB), short reads in *de novo* mode (SR_DN), mixed reads in reference-based mode (MR_RB), mixed reads in *de novo* mode (MR_DN), merged reads in *de novo* mode (MeR_DN), merged reads in reference-based mode (MeR_RB), and merged reads in both modes (MeR_RB_DN). Mixed reads refer to the combined use of short and long reads within a single assembly process, whereas merged reads indicate separate assemblies of short reads and mixed reads, followed by merging the resulting assemblies. For each approach, StringTie2 performance (Table S2) was quantified in terms of accuracy metrics, reflecting the ability to correctly identify transcript isoforms (Figure 1B). SR_DN approach achieved the lowest accuracy, ranging from 35.1% to 45.1% with its maximum at -c 1.5 and -f 0.02. Surprisingly, generally, the highest accuracy rate of 81.1% was obtained when predictions from both short and mixed reads in both modes were merged (MeR_RB_DN). This outcome underscores the synergistic benefits of integrating multiple read types and assembly strategies.

Consequently, our pipeline employs a dual-version strategy—by default, StringTie2 version 2.1.1 is used for assembling short-read data and version 2.2.1 for mixed-read data, ensuring optimal performance across a range of sequencing data types. Users should note that while these versions provided robust performance in our tests, alternative versions may be considered depending on specific experimental contexts and can be adjusted via command-line. We set -c 1.5 and -f 0.02 based on pilot experiments with SIRVs, which balanced sensitivity and precision in isoform detection (see Table S2). After completing individual assemblies, we incorporated the MeR_RB_DN approach to merge reference-based and de novo transcripts, resulting in a comprehensive and unified transcriptome annotation that captures a diverse array of transcript isoforms.

### Identification of artifactual transcripts

In our tests with *in-silico* data, we identified several types of artifactual transcripts, including those with unusually long exons exceeding 100,000 nt, reversed duplicates (transcripts overlapping the same genomic regions as the reference ones but on the opposite strand), as well as fragmented and chimeric RNA. Fragmented transcripts are individual RNAs that have been split into multiple predicted gene models due to incomplete transcript reconstruction or assembly errors. Chimeric transcripts, on the other hand, erroneously combine sequences from multiple distinct genes. In eukaryotic genomes, exons generally range in size from approximately 50 to 5,000 nt, with the majority falling between 100 and 1,000 nt [48]. Instances of exons exceeding 10,000 nt are exceptionally rare and are usually associated with specific organismal features, such as immune-related genes or tandem duplications. Even in these rare cases, such lengths remain an order of magnitude below 100,000 nt [49]. While highly effective in many contexts, StringTie2 may incorrectly model transcripts, especially in regions with overlapping genes or incomplete stop codons. Such inaccuracies can extend exon boundaries far beyond biologically realistic lengths [50].

To address the identified challenges and eliminate artifacts, we added a custom function to the pipeline that ensured that transcripts with exons exceeding 10,000 nt and/or transcripts that surpassed 100,000 nt of the genomic region were removed, thus refining the annotation and preventing the propagation of erroneous data. Additionally, we included a function to identify and filter out reversed duplicates without inadvertently removing legitimate transcripts. We also implemented criteria to flag potential fragmented and chimeric RNA based on exon length, genomic proximity, and functional annotations. In our pipeline, indicators for fragmentation include multiple predicted genes in close genomic proximity (within 1,000 nt) on the same strand and adjacent or overlapping genomic coordinates. The exon length and genomic proximity thresholds are based on general eukaryotic genome characteristics [49]. These parameters should be adjusted as needed based on the specific organism or dataset characteristics. Transcripts and genes are flagged as chimeric if they meet at least one of the following criteria: (i) they display structural inconsistencies, such as widely spaced or non-contiguous exons; (ii) they span multiple distinct genomic regions; or, in the case of genes, (iii) they possess multiple, unrelated functional annotations. As part of our automated pipeline, we only added information indicating whether a gene or transcript is possibly fragmented or chimeric. The decision to remove corresponding genes or transcripts should be made by the user based on specialized evaluation. Incorporating functional annotations into the pipeline as a part of manual curation can significantly improve the accuracy of detecting artifacts.

### Pipeline overview and workflow

Having resolved the challenges outlined above, we developed a versatile genome annotation pipeline and demonstrated its performance by applying it to the asexual strain of *S. mediterranea.* This pipeline integrates multiple sequencing technologies and bioinformatics tools to achieve comprehensive and accurate gene and transcript annotations across different organisms. The pipeline is divided into two main parts, each structured into distinct sequential steps.

#### Part I: Pre-processing and assembly (SmedAnno)

The first part of the pipeline is container-orchestrated, fully automated, and encompasses pre-processing, alignment, assembly, and functional annotation. The input data for the pipeline can be either short-read or mixed, the latter including both short and long reads. While even a single sample can be processed when only short reads are available, incorporating multiple samples is recommended to achieve more reliable and comprehensive results. Mixed-read datasets require at least one sample with short reads and one with long reads. Additionally, the automated pipeline is optimized for paired-end reads in short-read mode. Mapping and transcript assembly are performed independently for each sample. The workflow begins by pre-processing the RNA-seq data, performing adapter and quality trimming with TrimGalore and Filtlong for short- and long reads, respectively, followed by optional length and quality filtering. Next, the pipeline focuses in particular on the removal of reads corresponding to ribosomal RNA (rRNA), based on their alignment to user-provided rRNA reference sequences (Figure 2). This step was found to be very important in the case of organisms with unusually high A/T content, such as *S. mediterranea*. Library preparation methods typically involve rRNA depletion or polyA enrichment to facilitate the study of protein-coding gene expression. However, commercially available rRNA depletion kits are mostly optimized for model species, and polyA enrichment is not effective in organisms with a high A/T content. Consequently, reads corresponding to rRNA can be abundant in sequencing data for non-model organisms. In scenarios where rRNA reference sequences are unavailable (not provided by the user), the pipeline bypasses the rRNA removal step, continuing with the whole RNA-seq dataset. The pipeline then aligns the processed RNA-seq reads to the reference genome using STAR for short reads and Minimap2 for long reads. After completing this step, splice junctions can be optionally identified using a deep-learning-based predictor DeepSplice [41]. It uses convolutional neural network architectures (OpenSpliceAI-like) and supports CUDA acceleration for efficient computation. DeepSplice infers splicing donor and acceptor motifs directly from the reference genome, based on an integrated model. Currently the user can select a model, according to phylogenetic proximity to the target organism, from five available ones (human, mouse, zebrafish, honeybee, and thalecress). Next, the transcript assembly phase uses StringTie2 – by default, version 2.1.1 is applied to short-read data, while version 2.2.1 is used for mixed-read data (which integrates both short and long reads). Users can specify alternative versions if desired. After assembly, the pipeline merges transcripts across all samples using StringTie2 to create unified sets of transcripts. This merging process is performed separately for reference-based (if reference annotation was provided) and *de novo* assemblies before combining them into the final complete annotation. This dual approach by performing both reference-based and *de novo* annotation, provides comprehensive assembly, thus capturing a wide range of transcript isoforms. DeepSplice evidence is then merged with StringTie2 predictions. Subsequently, transcripts with excessively long exons or that cover unusually large genomic regions are filtered out. Next, the pipeline uses gffcompare to compare the predicted annotations against the reference annotations (if provided), assessing the completeness and accuracy of the assembled transcripts. This comparison generates a report that includes data summaries and accuracy estimations. Additionally, it enriches the predicted annotations with detailed information about their overlaps with known reference transcripts and genes. However, the generated annotations may contain issues that can compromise the quality and usability of the annotation files. To address these issues, the pipeline employes AGAT (Another GTF/GFF Tool), which systematically identifies and corrects common syntax errors, ensuring that essential attributes such as gene_id and transcript_id are present for each feature, preventing incomplete annotations, correcting improper use of delimiters in attribute fields (such as replacing spaces with a semicolon), detecting and removing duplicate entries, and ensuring correct key-value pair syntax. This correction phase is crucial for maintaining high-quality genome annotation. The obtained set of transcripts is then subjected to functional annotation in a process that involves the sequential application of several tools, and which includes homology analysis and the identification of protein domains. To this end, first, the open reading frames (ORFs) are predicted using TransDecoder. Following that, BLASTp and BLASTx are used for homology search through the SwissProt database. Finally, protein domains are identified using hmmscan through the PFAM database and InterProScan, which aggregates multiple domain resources (SMART, Panther, TIGRFAM, etc.) and enriches predictions with Gene Ontology (GO). This multi-faceted approach filters out low-confidence annotations, retaining only the transcripts supported by multiple lines of evidence. We used the SwissProt, PFAM and InterProScan databases as a part of the automated pipeline specifically for their low to medium memory requirements. However, we recommend to use additional resources, for example eggNOG, NCBI, UniRef, KEGG, RefSeq, COG, Reactome, and BioCyc for conducting in-depth homology analyses, as these databases provide complementing information, evolutionary relationships, and diverse protein characteristics. Next, the functional annotations are integrated into the GTF file. This integration applies default thresholds: for PFAM predictions, a bit score greater than 20, a conditional e-value below 1e-5, and coverage exceeding 50%; for SwissProt predictions, a BLAST e-value threshold of 1e-5 and an identity threshold of 25%; InterPro hits are retained if they meet the e-value threshold of < 1e-5 in at least one of the member databases. These criteria ensure that low-confidence functional annotations are discarded. However, all filtering thresholds can be configured by the user. Then, the pipeline adds new columns to the GTF annotation file, including has_ORF (TRUE/FALSE), best_SwissProt_blastp_hit, best_SwissProt_blastx_hit, best_Pfam_hit, and best_InterPro_hit domain information. The next step is the identification of overlapping genes and transcripts, which detects genes that share genomic regions and checks if each transcript in a gene has at least one overlapping exonic region with any other transcript in the same gene. Lastly, the pipeline identifies the aforementioned reversed duplicates and addresses fragmented and chimeric genes and transcripts. At the completion of functional annotation, the pipeline emits an interactive summary-and-advice block. Six boolean quality control flags (*overlapped_predicted_gene*, *is_potentially_fragmented_reference, is_potentially_chimeric_reference, is_potentially_fragmented_de_novo, is_potentially_chimeric_structure_de_novo, and is_potentially_chimeric_function_de_novo*) are printed together with plain-language recommendations (e.g. “merge fragmented loci”, “split structural chimeras”). Users are expected to review each flagged locus individually in IGV, confirm read support, and decide case-by-case whether to merge, split, or retain the model unchanged. All these steps (Figure 2) are implemented in the SmedAnno tool, forming Part I of the entire pipeline. The source code can be accessed on GitHub (https://github.com/Norreanea/SmedAnno).

**Figure 2.**
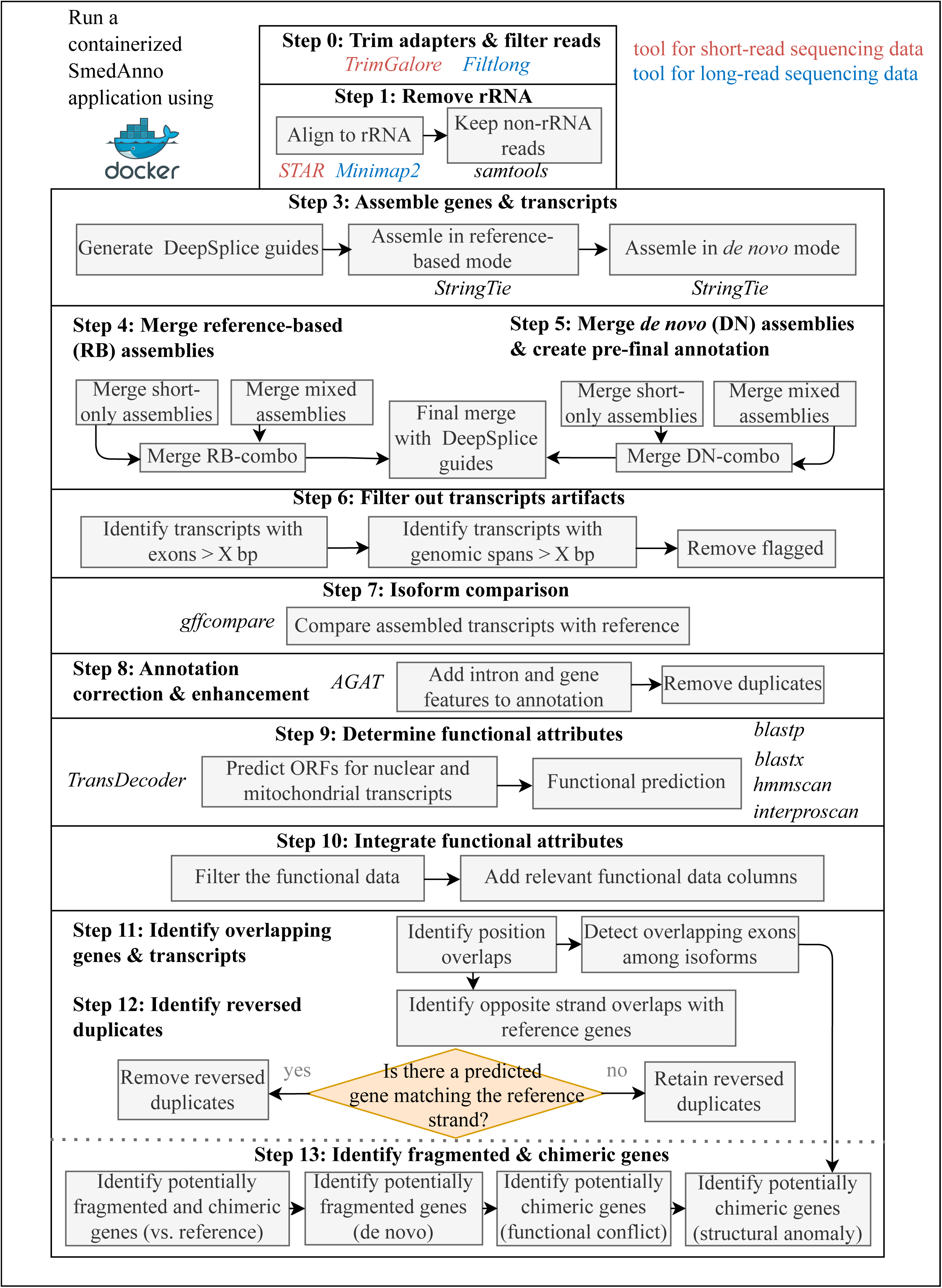
Workflow diagram for RNA-seq pre-processing, transcript assembly, and functional annotation.

#### Part II: Manual curation and refinement

The second part of the pipeline involves user-driven inspection and refinement of the annotated data. Despite the implementation of several automated steps aimed at optimizing transcript assembly and obtaining biologically relevant transcripts that encode functional protein domains, resolving overlapping or fragmented transcripts remains a significant challenge that requires specialized approaches. While the primary focus is on transcript assembly, gene features are subsequently derived from the assembled transcripts. Fragmented and chimeric transcripts, which primarily affect RNA-level annotations, are addressed by prioritizing the longest isoform to define each gene, thereby reducing redundancy and improving annotation accuracy. To accurately determine whether genes should be merged after being marked as potentially fragmented and/or overlapped, promoter and terminator sequences should be analyzed using Promoter 2.0 and ARNold tools (Figure 3). This analysis verifies the coherence of regulatory regions, ensuring that merged gene models maintain consistent promoter and terminator elements. These tools are accessible via their respective web pages and are not included in the automated pipeline. Users must extract sequences between inspected genes to check for the presence of promoters and terminators. After potential merges, manual adjustments are advised to finalize gene models, ensuring accurate gene length prediction and overall annotation reliability.

**Figure 3.**
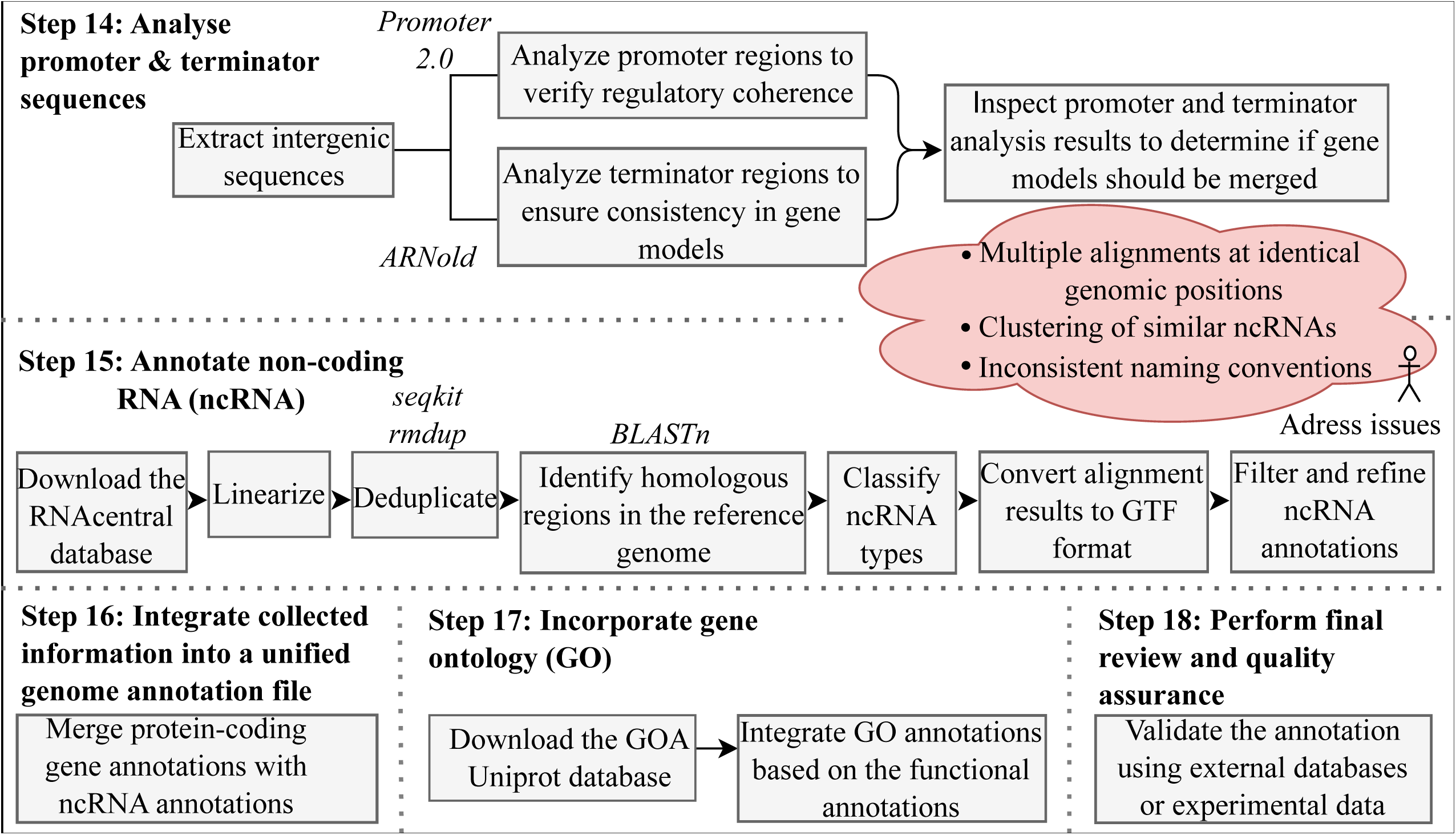
Workflow diagram for user-driven inspection and refinement.

While genome annotation typically focuses on protein-coding genes, our pipeline also includes a step for annotating non-coding RNAs (ncRNAs), which have significant functional relevance. This step is optional, as it requires downloading the extensive (>20 GB) RNAcentral database. The ncRNA sequences are first linearized and deduplicated using seqkit rmdup to remove redundant entries. Homologous regions are then identified in the reference genome using BLASTn. After the identification step, ncRNA types are classified based on the sequence identifier. The matching results are then converted to GTF format. A multi-step filtering and refinement process is used to improve the accuracy of the annotations. This process addresses several common issues, including multiple alignments at identical genomic positions, inconsistent naming conventions, and clustering.

The final step integrates all gathered information into a unified genome annotation file, ready for downstream applications, including both coding and non-coding elements with comprehensive functional insights. The bioinformatics tools used in each pipeline step, along with their respective versions are summarized in Table 1.

**Table 1.** Bioinformatics tools and pipeline steps.

| Step | Tool | Version |
| --- | --- | --- |
| Step 0: Trim adapters & filter reads | TrimGalore<br>Filtlong | 0.7.1<br>0.2.1 |
| Step 1: Remove rRNA | STAR<br>Minimap2<br>samtools | 2.7.9a |
| Step 2: Align reads to the reference genome | STAR<br>Minimap2 | 2.7.9a |
| Step 3: Assemble genes & transcripts | StringTie2<br>DeepSplice | 2.1.1 (short reads)<br>2.2.1 (mixed reads)<br>0.10 |
| Step 4: Merge reference-based assemblies | StringTie2 | 2.2.1 |
| Step 5: Merge *de novo* assemblies & create pre-final annotation | StringTie2 | 2.2.1 |
| Step 6: Filter out transcripts artifacts | bash |  |
| Step 7: Compare isoforms | gffcompare | 0.12.6 |
| Step 8: Correct & enhance annotation | AGAT | 1.4.1 |
| Step 9: Determine functional attributes | TransDecoder<br>blastp<br>blastx<br>hmmscan<br>InterProScan | 5.7.1<br>2.15.0+<br>2.15.0+<br>3.3.2<br>5.67 |
| Step 10: Integrate functional attributes | R | 4.4.1 |
| Step 11: Identify overlapping genes & transcripts | R | 4.4.1 |
| Step 12: Identify reversed duplicates | R | 4.4.1 |
| Step 13: Identify fragmented & chimeric genes | R | 4.4.1 |
| Step 14: Analyse promoter & terminator sequences | Promoter 2.0<br>ARNold | No official version<br>number<br>No official version<br>number |
| Step 15: Annotate non-coding RNA (ncRNA) | blastn<br>seqkit | 2.15.1+<br>2.8.2 |
| Step 16: Integrate collected information into a unified genome annotation file | R | 4.4.1 |
| Step 17: Incorporate gene ontology (GO) | R | 4.4.1 |
| Step 18: Perform final review and quality assurance | R | 4.4.1 |

### SmedAnno benchmarking

To benchmark the automated Part I of our pipeline (SmedAnno), we compared its performance against BRAKER3, a widely used genome annotation tool that integrates GeneMark-ET and AUGUSTUS gene prediction, guided by short-read RNA-seq data [39]. The comparison was carried out using three genomes: two classic models – *A. thaliana* (AT) and *C. elegans* (CE) – and *S. mediterranea* (SM). Since BRAKER3 utilizes *ab initio* approach and short-read data only, we ran a scaled-down version of SmedAnno without reference-based guidance, exclusively on Illumina reads, to ensure a valid comparison. For *A. thaliana* and *C. elegans*, publicly available data were used, while for *S. mediterranea* we employed in-house generated datasets (Table S1). For DeepSplice, which is integrated into our pipeline, we employed thalecress model for *A. thaliana*, and honebee model for *C. elegans* and *S. mediterranea*. SmedAnno was run in four parameter configurations (with/without DeepSplice and TransDecoder). After assembling transcripts for each genome, we evaluated the resulting transcriptomes using BUSCO, which assesses the presence of conserved orthologs, by comparing a transcriptome of interest to those within the Metazoa lineage, and gffcompare.

BRAKER3 reached slightly higher BUSCO completeness in AT (98.9 %) and CE (98.6 %) than the most comparable SmedAnno runs (∼95 % and 88.8 %, respectively), but fell behind in SM (76.9 % vs 78.6 % for both DeepSplice –TD and noGuide –TD) (Figure 4A). Notably, SmedAnno yielded increased fraction of duplicated BUSCOs for all tested genomes (Figure 4A). We found that SmedAnno configurations without TransDecoder generated more transcripts being identified as single-copy BUSCOs. Next, the annotations obtained with BRAKER3 and SmedAnno were compared with corresponding reference genome annotations using gffcompare. The proportion of true positive predictions was assessed at transcript, exon and intron levels, and depicted as F1 scores, where the higher value better reflects the reference. As expected, BRAKER3 performed better in CE. BRAKER3’s advantage in *C. elegans* is consistent with its AUGUSTUS species-specific parameter set, which was trained on that very genome. SmedAnno+DeepSplice (-TD) outperformed BRAKER3 in tests on AT and SM, displaying higher F1 score at the exon and transcript levels in AT, and at all levels in SM (Figure 4B).

**Figure 4.**
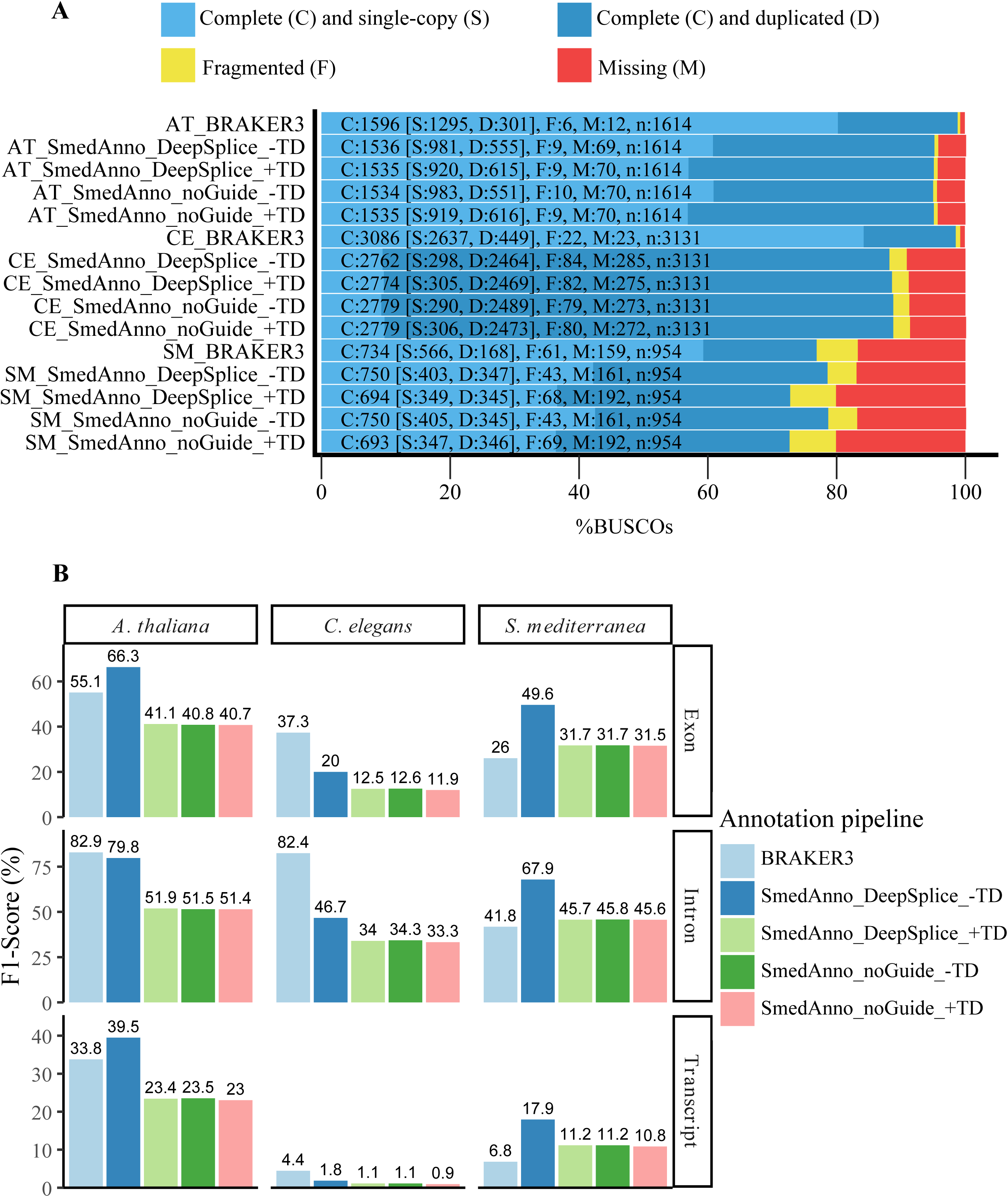
Benchmark of SmedAnno versus BRAKER3 on short-read data. (A) - BUSCO completeness for annotations of *A. thaliana* (AT), *C. elegans* (CE) and *S. mediterranea* (SM) transcriptomes. Stacked bars depict the proportion of: complete single-copy, complete duplicated, fragmented and missing BUSCOs for each pipeline: BRAKER3; SmedAnno with DeepSplice junction hints merged but without TransDecoder (SmedAnno_DeepSplice_-TD); the same plus TransDecoder (+TD); and SmedAnno runs without DeepSplice guidance (noGuide_±TD). Numbers inside the bars correspond to raw BUSCO counts. (B) - Structural accuracy. Bars show F1 scores (%, harmonic mean of sensitivity and precision) at exon, intron and transcript level for the same pipelines. Colours denote pipelines, panels group the three species.

### Genome annotation of the asexual strain of *S. mediterranea*

After demonstrating the efficiency of SmedAnno, we applied the whole pipeline to generate a genome annotation of the asexual strain of *S. mediterranea*. To this end, we used the available reference genome annotation and both Illumina and PacBio sequencing datasets (Table S1). The SmedAnno output was subjected to manual curation and refinement, which included identification of artifactual transcripts and detection of missing genes. The annotation, schMed_Pn, is available in GEO repository under accession GSE294569.

### Identifying fragmented and chimeric artifacts

In the context of genomic annotation, artifacts are erroneous gene or transcript models that arise from limitations in the annotation process rather than representing true biological entities. Specifically, we refer here to fragmented artifacts as those that occur when a single gene is incorrectly split into multiple separate gene models. This fragmentation can result from incomplete transcript assembly or sequencing errors, leading to discontinuous or truncated representations of what is biologically a single gene. On the other hand, chimerism typically arises from misassembly during transcriptome reconstruction, especially in regions with high gene density or overlapping gene structures.

SmedAnno employs flags such as: *is_potentially_fragmented_de_novo*, *is_potentially_chimeric_structure_de_novo*, and *is_potentially_chimeric_function_de_novo* to identify these artifacts based on functional and structural criteria. Below, we discuss three illustrative examples from schMed_Pn (depicted in Figure 5) that highlight the complexities and implications of these annotation challenges. SMESGs here are reference genes and MSTRGs are putative genes.

**Figure 5.**
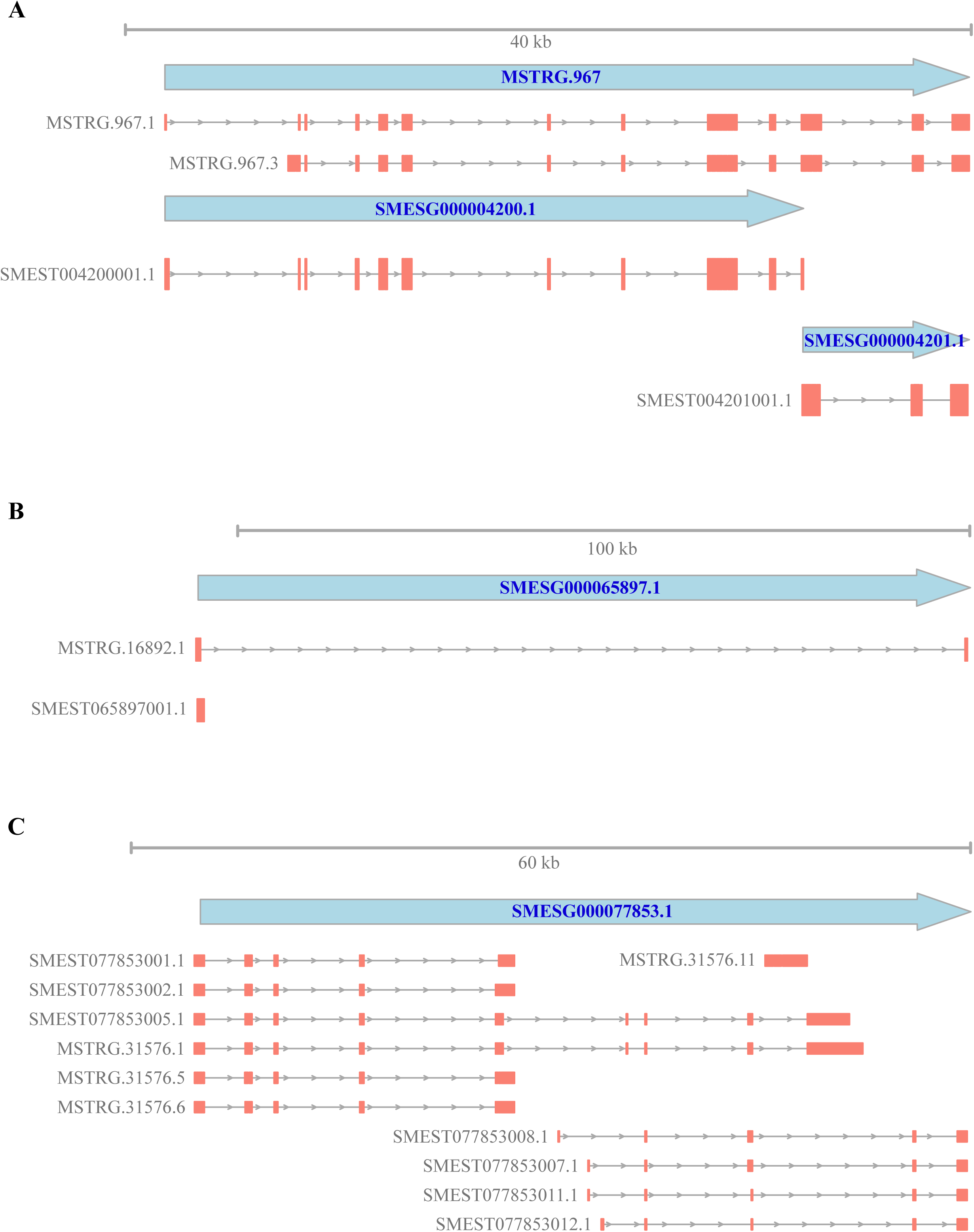
Representative annotation artifacts generated by SmedAnno pipeline. (A) - Fragmented gene. (B) Chimeric transcript. (C) - Functionally chimeric gene with divergent transcript annotations. Genes are depicted with cyan arrows, while transcripts are shown as combinations of gray lines and orange rectangles, denoting introns and exons, respectively.

#### Prediction of gene fragments

In one genomic region, three genes—SMESG000004200.1, SMESG000004201.1, and MSTRG.967—were located in close proximity on chromosome 1 (smed_chr1) (Figure 5A). Specifically, gene SMESG000004200.1 spanned position 14,973,263 to 15,003,452 on the positive strand, while SMESG000004201.1 extended from 15,003,453 to 15,011,292 on the same strand. StringTie2, predicted an additional transcript, MSTRG.967, which encompassed the regions of both genes (14,973,256 to 15,011,292). Both SMESG000004200.1 and SMESG000004201.1 were independently annotated as “Epidermal growth factor receptors.” However, the prediction by StringTie2 that merged these two genes into a single transcript was more biologically plausible. This is because the close genomic proximity and shared functional annotations suggested a possible complex gene structure that is better represented by a single, contiguous transcript. Flagging these genes as potentially fragmented (*is_potentially_fragmented_de_novo*) highlighted the necessity to consider merged annotations in densely packed genomic regions to avoid misinterpretation of gene boundaries.

#### Chimeric transcript prediction due to excessive intron length

The gene SMESG000065897.1 initially comprised a single transcript, SMEST065897001.1, spanning positions 325,241,809 to 325,242,781 on chromosome 1 (Figure 5B). However, StringTie2 predicted an additional transcript, MSTRG.16892.1, which extended dramatically from 325,241,809 to 325,347,749, introducing an intron exceeding 100,000 nt. This unusually large intron length is atypical and suggested a misassembly, classifying MSTRG.16892.1 as potentially chimeric (*is_potentially_chimeric_structure_de_novo*).

The presence of such a large intron likely indicated that MSTRG.16892.1 erroneously combined sequences from multiple genomic loci or failed to accurately represent the true transcript structure of SMESG000065897.1. Consequently, the chimeric transcript should be disregarded, and the gene localization should remain unchanged, basing solely on the original transcript SMEST065897001.1.

#### Functionally chimeric gene due to divergent transcript annotations

The gene SMESG000077853.1 exhibited a complex transcriptional landscape (Figure 5C). While most transcripts, such as MSTRG.31576.1, MSTRG.31576.5, MSTRG.31576.6, and all reference SMEST transcripts were annotated as “ankyrin repeat proteins,” one transcript, MSTRG.31576.11, was annotated as “Squamous cell carcinoma antigen recognized by T-cells 3 (SART-3)”. This divergence in functional annotation across transcripts resulted in the gene being flagged as chimeric (*is_potentially_chimeric_function_de_novo*). These inconsistent functional annotations suggested that MSTRG.31576.11 may have been incorrectly assembled, possibly incorporating sequences from a different gene or representing a mispredicted splice variant. Given that “SART-3” functions are unrelated to “ankyrin repeat proteins,” it is evident that MSTRG.31576.11 does not belong to SMESG000077853.1. Consequently, MSTRG.31576.11 was filtered out.

In total, the automated run flagged 5.1% of all gene models, of which 1.2% were confirmed as artifacts upon manual inspection.

### Detecting missing genes through functional annotation and homology assignments

Functional annotations and homology assignments were carried out to explain the biological function of the assembled transcripts and identify orthologous genes by using established databases and computational tools. This approach facilitated the detection of missing genes, thus highlighting potential gaps and guiding further refinement of the transcriptome assembly.

To incorporate functional data into the newly developed *S. mediterranea* genome annotation (schMed_Pn), we used the following databases: Pfam, SwissProt, and non-redundant (nr) protein sequences from NCBI. Our annotation comprised 25,265 genes, of which 4,011 were novel and 21,254 were identical to the reference genes. Each gene possessed at least one function-related entry, either through a database hit or by being classified as having an ORF. For 18,552 genes, we successfully identified Gene Ontology (GO) terms using homology-based strategies, which included the pfam2go package, UniProt GOA conversion, and a custom R script (Table S3) to retrieve GO annotations based on NCBI IDs. Notably, 2,427 of these GO-annotated genes were putative. To evaluate the similarity of GO terms between novel and reference genes, Venn diagrams were constructed, highlighting shared and unique functional categories across the gene sets. We assigned a total of 23,909 GO terms to genes within our annotation. Specifically, 18,586 GO terms were assigned to SMESG (reference) genes, and 5,323 GO terms were assigned to MSTRG (putative) genes. Of these, 5,180 GO terms were shared between both gene categories, while 143 GO terms were uniquely assigned to MSTRG genes (Figure 6A). To visualize the functional distribution of the gene sets, we generated word clouds representing the most frequent unique GO terms for each group: MSTRG (Figure 6D) and SMESG (Figure 6B), as well as those that were common for two groups (Figure 6C).

**Figure 6.**
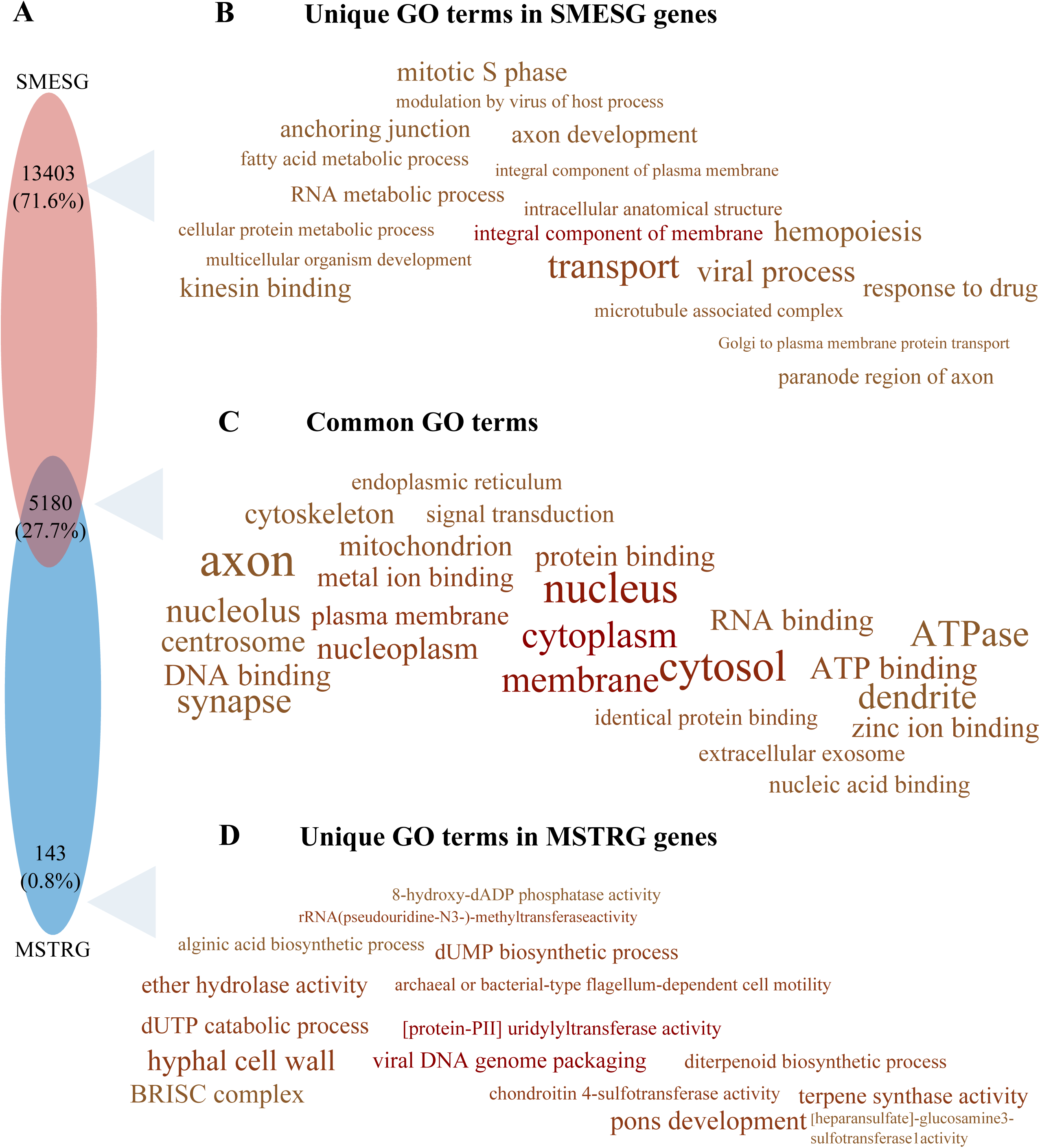
Venn diagram and a word cloud of GO terms assigned for novel and reference genes for the *S. mediterranea* asexual strain. (A) - Venn diagram of GO term overlaps between MSTRG (novel genes) and SMESG (reference genes). (B) - Word cloud of unique GO terms assigned to SMESG (reference) genes. (C) - Word cloud of common GO terms assigned to both SMESG and MSTRG genes. (D) - Word cloud of unique GO terms assigned to MSTRG (novel) genes.

### Insights into planarian ncRNA diversity

Beyond refining the annotations of protein-coding genes, we also performed a preliminary assessment of non-coding RNAs (ncRNAs) in *S. mediterranea*. Although not an exhaustive characterization, our analysis revealed a diverse repertoire of regulatory RNAs, highlighting significant differences between publicly available annotations and newly generated data. Publicly available datasets from RNAcentral assigned a total of 154 rRNAs, 578 miRNAs (including 269 precursors), 61 tRNAs, 2 snoRNAs, 12 snRNAs, and 2 SRP RNA hits to *S. mediterranea*. After deduplication, these numbers were refined to 129 rRNA, 571 miRNAs (with 156 precursors), 60 tRNAs, 2 snoRNAs, 12 snRNAs, and 2 SRP RNA hits. Additionally, according to recent literature [47], *Schmidtea* genome contains 457 tRNA hits, which were reduced to 434 after deduplication. In contrast, using our annotation pipeline, we identified 154 rRNA hits across 2,287 genomic positions, 179 miRNA hits across 1,042 genomic positions, 488 tRNA hits distributed over 4,298 genomic positions, 2 snoRNA hits across 69 genomic positions, 18 snRNA hits across 141 genomic positions, 6 miscellaneous RNAs spread across 4,752 genomic positions, 18 RNase P ncRNA components across 408 genomic positions, 21 lncRNA hits across 185 genomic positions and 2 sRNA hits across 187 genomic positions. The discrepancy between published records assigned to *S. mediterranea* and our annotated RNAs highlights the ongoing need for careful re-annotation and experimental validation in underexplored organisms like planarians. In addition, we uncovered 40 copies of regions encoding signal recognition particle RNAs (SRP_RNAs), localized on chromosome 3 and flanked by the genes SMESG000020110.1 (BLAST-predicted as a lysine-specific demethylase) and MSTRG.33721 (BLAST-predicted as containing a thrombospondin type 3 repeat). Moreover, we identified 10 copies of TERC_RNA located on smed_chr4, while *TERT* gene (SMESG000061912.1) was located on smed_chr3.

### Comparative analysis of selected *S. mediterranea* genome annotations

To ensure the reliability of schMed_Pn genome annotation, we compared it with commonly used genome annotations of *S. mediterranea*. First, we analyzed transcript length distributions. It revealed that all annotations predominantly consisted of relatively short transcripts, as reflected by a sharp decline in cumulative transcript counts with increasing transcript length (Figure 7A). Specifically, dd_Smed_v6 showed the steepest decline in cumulative transcript counts with increasing length, suggesting a higher fragmentation rate; however, caution is advised in interpreting these distributions, as there is no “standard” shape for planarian transcript length. In contrast, schMed_Pn demonstrated a more gradual decline, which indicated a broader distribution of transcript lengths. Assemblies such as schMedS1, schMedS2, schMedS3_1, and schMedS3_2 exhibited moderate slopes with slight differences, indicating varying transcript length distributions among them. The SmedAsxl genome_v1.1 displayed a narrower range of longer transcripts. In contrast, SmedGD_c1 and SMESG.1, similar to schMed_Pn, demonstrated relatively smooth distributions, signifying balanced transcript coverage with a significant proportion of longer transcripts.

**Figure 7.**
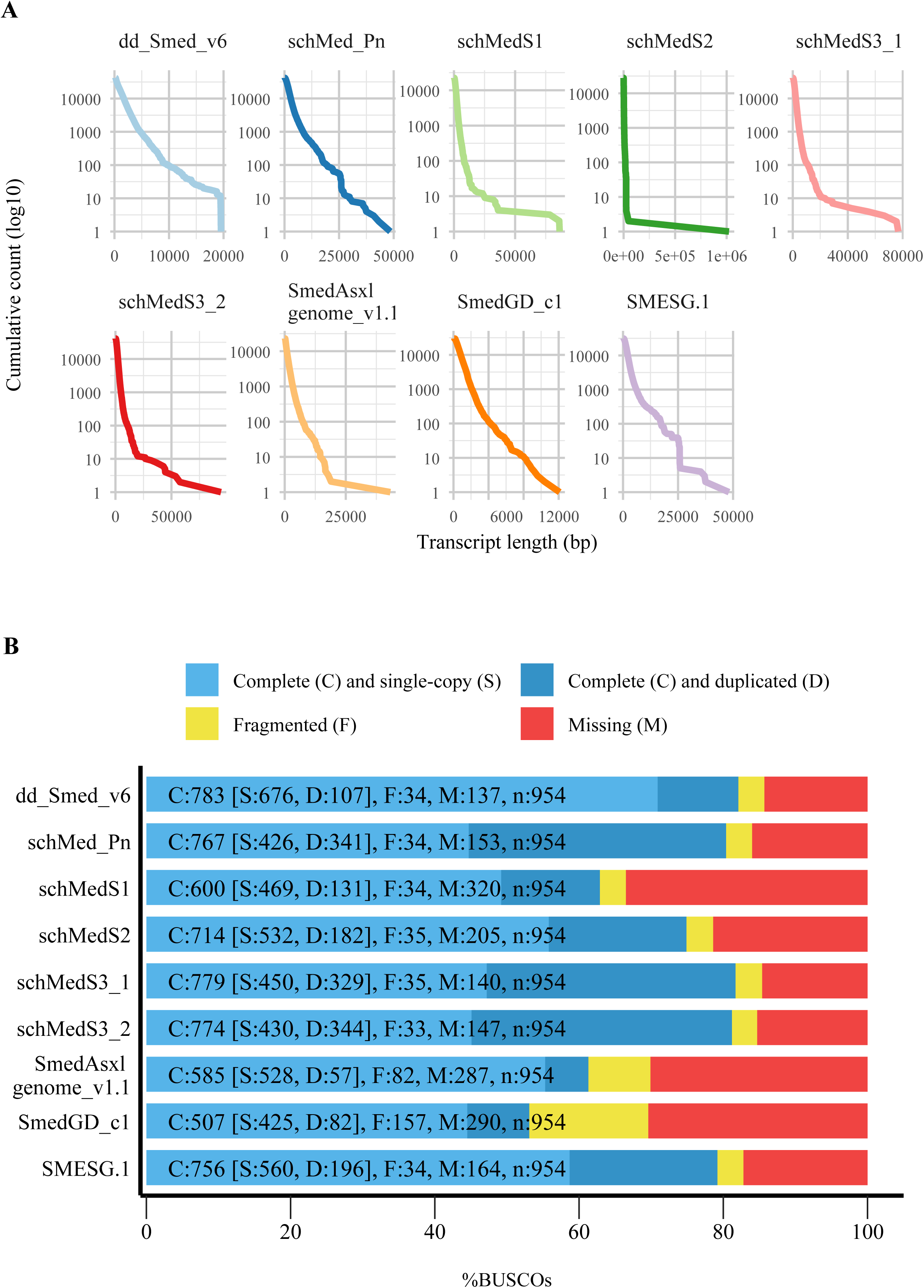
Comparative analysis of selected *S. mediterranea* genome annotations. (A) - Cumulative distribution of transcript lengths in selected annotations. The y-axis (logarithmic scale) represents the cumulative count of transcripts/isoforms, indicating how many transcripts exceed each length threshold on the x-axis (linear scale). (B) - BUSCO completeness of selected *S. mediterranea* annotations.

The completeness of the transcriptome assemblies was evaluated using BUSCO (Figure 7B). The dd_Smed_v6 and SMESG.1 annotations achieved the highest single-copy ortholog recovery—783 and 756 complete BUSCOs out of 954, respectively—with low duplication rates (107 and 196 duplicated BUSCOs), indicative of very accurate core gene models but at the cost of fewer total transcript isoforms. In contrast, schMed_Pn attained a balanced profile of completeness and isoform richness, with 767 complete BUSCOs (426 single-copy, 341 duplicated), coupled with minimal fragmentation (34) and moderate missing counts (153). This reflected our pipeline’s emphasis on maximizing isoform discovery without substantially sacrificing core gene completeness. Other assemblies (schMedS1–3, SmedAsxl genome_v1.1) showed lower complete counts and higher fragmentation or missing rates, consistent with greater redundancy or gaps. Based on BUSCO analysis, both SMESG.1 and dd_Smed_v6 demonstrated robust performance (albeit with different trade-offs in duplication and missing orthologs). Notably, schMed_Pn also ranked among the stronger annotations, demonstrating moderate duplication and missing ortholog levels, while retaining high completeness.

In addition, we employed rnaQUAST and Transrate to generate detailed assembly quality metrics (Table 2). Notably, schMed_Pn exhibited a high number of transcripts over 1,000 nt and maintained a competitive N50 value (the length of the shortest contig or scaffold covering 50% of the total assembly), reflecting effective transcript coverage and solid assembly quality.

**Table 2.** Comparative metrics for selected transcriptome assemblies of *S. mediterranea*.

| *Metric* | dd_Smed_v6 | schMed_Pn | schMedS1 | schMedS2 | schMedS3_1 | schMedS3_2 | SmedAsxl_v1.1 | SmedGD_c1.3 | SMESG.1 |
| --- | --- | --- | --- | --- | --- | --- | --- | --- | --- |
| *No. of Transcripts* | 41,745 | 41,396 | 22,348 | 27,790 | 41,358 | 41,036 | 22,855 | 29,850 | 31,102 |
| *Min transcript length (nt)* | 201 | 200 | 267 | 267 | 145 | 131 | 75 | 87 | 267 |
| *Max transcript length (nt)* | 19,524 | 48,466 | 86,373 | 1,022,601 | 76,733 | 94,201 | 43,226 | 12,11 | 48,121 |
| ***Total Bases (M)*** | 53.23 | 97.09 | 40.12 | 57.93 | 72.60 | 71.81 | 25.55 | 23.15 | 65.55 |
| ***Mean transcript length (nt)*** | 1,275 | 2,345 | 1,795 | 2,084 | 1,755 | 1,750 | 1,114 | 770 | 2,108 |
| ***No. of transcripts >1k*** | 18,487 | 31,199 | 16,082 | 20,727 | 29,402 | 28,932 | 8,610 | 7,218 | 23,648 |
| ***No. of transcripts >10k*** | 94 | 702 | 67 | 293 | 116 | 119 | 47 | 2 | 346 |
| ***No. of transcripts with ORF*** | 20,461 | 32,798 | 18,694 | 23,854 | 31,503 | 31,147 | 12,937 | 17,567 | 26,665 |
| ***N50 (nt)*** | 1,931 | 3,070 | 2,159 | 2,535 | 2,133 | 2,141 | 1,653 | 1,011 | 2,570 |
| ***GC (%)*** | 0.34 | 0.33 | 0.33 | 0.33 | 0.33 | 0.33 | 0.34 | 0.36 | 0.33 |

Performance benchmarking enabled the identification of improvements and areas needing enhancement, while also positioning schMed_Pn within the context of current resources. These analyses collectively demonstrated that our approach offers a balanced and complete annotation, effectively capturing a wide range of transcript lengths and maintaining high completeness.

## Discussion

Numerous pipelines and tools have been developed to tackle genome annotation across diverse organisms. For well-studied species—such as *Drosophila melanogaster* or *Arabidopsis thaliana*—community-maintained platforms (FlyBase [51] and TAIR [52,53] respectively) provide robust resources that streamline annotation. In prokaryotes, automated tools like Prokka [10] efficiently handle smaller, less complex genomes, while eukaryotic model organisms often benefit from RefSeq [7] or large consortia efforts, which incorporate extensive experimental validation. However, non-model eukaryotes, including many invertebrates and plants, typically face an incomplete reference, novel repetitive elements, and limited training data for *ab initio* predictions. Pipelines such as MAKER [5,54], BRAKER [55,56], GGA [9], nf-core GenomeAnnotator [12], and others can automate workflow portions—especially when reference data or orthologous gene sets are available. However, in certain species with high genomic complexity or less functional annotation data, these tools may still underperform or require extensive manual curation.

In this study, we developed and validated a comprehensive genome annotation pipeline designed to address the challenges of working with non-classical model organisms, while generating high-quality annotations applicable to broader genomic research. Our pipeline uniquely combines: (i) the integration of multiple data types, (ii) automated artifact detection, (iii) dual approach assembly —reference-based plus *de novo*, and (iv) broad functional integration.

The BUSCO completeness of SmedAnno (the automated part of the pipeline) was within 4% of BRAKER3 in *A. thaliana*, surpassed it by approximately 2% in *S. mediterranea*, and lagged behind by approximately 10% in *C. elegans*. Notably, SmedAnno runs consistently doubled the proportion of duplicated BUSCO genes. Because BUSCO counts every alternatively spliced isoform as a duplicate, such inflation is widely regarded as a proxy for improved isoform recovery rather than assembly noise [57]. In practical terms, this means that the apparent redundancy in SmedAnno is biologically informative when transcript diversity, rather than the most parsimonious gene count, is the primary research goal.

Leveraging this enhanced isoform recovery, our analysis identified 4,011 novel genes and multiple previously uncharacterized isoforms and ncRNAs in the asexual strain, enriching the genomic landscape of *S. mediterranea*. Notably, the GO term analysis of putative MSTRG genes revealed enrichment in diverse biological functions. The occurrence of these genes necessitates further investigation to better understand their implications and potential significance. While our transcriptomic data offer strain-specific insights, it should be considered that the annotation relies on the genome of the sexual strain as the reference. Consequently, differences between strains, such as chromosomal translocation and other genetic variations, may compromise the accuracy of gene prediction and transcript assembly, leading to potential inconsistencies in annotation quality.

The existing, planarian-specific repositories—such as PlanMine [21], PLANAtools [58], WormBase [59], PlanoSphere [60], PlanNET [61], and various planaria single-cell datasets [62–65]—provide essential resources for gene model comparisons and functional information in *S. mediterranea*. Despite their value, discrepancies in naming conventions, annotation updates, and differences between sexual and asexual strains can lead to inconsistencies. These challenges highlight the importance of robust, well-documented pipelines that can reconcile or supplement these external databases, thus ensuring more accurate gene models.

A detailed inspection of the curated annotation uncovered two intriguing features of the genomic architecture: genes located in close proximity and non-contiguous transcripts. Although these observations could potentially represent annotation artifacts, it is crucial to carefully assess them in the context of established genomic knowledge. Genes located in close proximity can reflect various biological scenarios. One possibility is the presence of functionally independent genes that happen to be closely spaced, as observed in other genomes where gene density is high [66,67]. For example, two planarian genes—SMESG000043996.1 and SMESG000043998.1— that encode kynureninase and synaptotagmin XVII, respectively, are unrelated but positioned close to each other. Alternatively, such arrangements may arise from evolutionary events such as gene duplication followed by functional divergence, where both copies remain in close physical proximity. Another scenario includes operon-like gene structures commonly seen in prokaryotes but also identified in some eukaryotic contexts [68]. These structures allow the co-regulation of adjacent genes through shared promoters or other regulatory elements. Bidirectional promoters, enabling two genes to be transcribed in opposite directions from the same regulatory region, represent another mechanism driving close gene arrangements [69]. On the other hand non-contiguous transcripts, where exons are unusually spaced or appear fragmented, often arise from structural variations, alternative splicing, or sequencing artifacts. Biologically, large introns or dispersed exonic regions can be linked to long-range regulatory elements or structural genome features such as transposon insertions. Moreover, such patterns may indicate complex gene models, as seen in genes encoding for proteins with repetitive domains [70,71] or in cases where alternative promoters and polyadenylation sites lead to diverse transcript forms [72,73]. Consequently, accurately capturing these intricate gene architectures often requires more comprehensive strategies than those employed for simpler genomes, underscoring the need for user scrutiny and flexibility in annotation pipelines. The primary advantage of our pipeline lies in its capacity to address these complexities. By combining reference-based and *de novo* methods and incorporating both coding and non-coding RNA annotations it addresses the challenges posed by scarcely studied genomes. However, the pipeline’s reliance on user-defined thresholds and manual curation introduces subjectivity, potentially impacting reproducibility.

While the automated steps of SmedAnno effectively identify fragmented and chimeric transcripts, resolving these artifacts requires expertise and thus is not fully automated. Another limitation concerns functional annotations, which can lead to false positives in identifying chimeric genes. For instance, the same protein from different species or proteins representing different subunits of a complex, may be annotated as distinct entries in functional databases. This can lead to genes being incorrectly flagged as chimeric if the functional entries do not exactly match. Accordingly, development of refined algorithms that account for near-identical annotations and minor terminological differences is vital.

Future enhancements could include the integration of machine learning algorithms to automate artifact resolution and improve annotation consistency. The present container infrastructure already simplifies the implementation of graphics processing units (GPUs) and central processing units (CPUs). Expanding the pipeline to support multi-omics data integration, such as proteomics and epigenomics, would further enhance its utility. Community-driven efforts to standardize annotation protocols and databases would ensure broader applicability and reproducibility. Finally, experimental validation of novel genes and isoforms identified through this pipeline will be critical to confirm their biological relevance.

Compared to pipelines that excel in either prokaryotic or model vertebrate annotation – where reference data is abundant and well-curated – our approach fills the gap for non-classical model systems with more limited resources. In doing so, it advances genome annotation efforts for *S. mediterranea* and provides a robust framework for addressing similar challenges in other species, fostering broader genomic discoveries.

## Availability of Source Code and Requirements

Project name: SmedAnno

Project homepage: https://github.com/Norreanea/SmedAnno

Operating system: Linux

Container image: Docker ≥20.10, docker-compose ≥2.5

Programming languages: shell, R, Python, awk

License: MIT License

## Data Availability

The *in-silico* datasets supporting the results and all other code used in this article are available in the GitHub repository archived in Zenodo https://doi.org/10.5281/zenodo.15130394. RNA-seq data can be accessed from the following GEO repositories: GSE293999 and GSE293997 from NCBI BioProject PRJNA1243063; and GSE294569 from NCBI BioProject PRJNA1246370. The final genome annotation file is available in GSE294569.

## List of Abbreviations

BP: base pairs
NT: nucleotide
RNA: ribonucleic acid
rRNA: ribosomal RNA
ncRNA: non-coding RNA
SIRV: Spike-In RNA Variants
ORF: open reading frame
GTF: Gene Transfer Format
BLAST: Basic Local Alignment Search Tool
STAR: Spliced Transcripts Alignment to a Reference
BUSCO: Benchmarking Universal Single-Copy Orthologs
GO: Gene Ontology
cDNA: complementary DNA
DNA: deoxyribonucleic acid
RIN: RNA Integrity Number
GGA: Galaxy Genome Annotation

## Declaration of generative AI and AI-assisted technologies in the writing process

During the preparation of this work, the authors used the free version of OpenAI-ChatGPT c2022 (v.GPT-o3-mini, February 2025) in order to improve manuscript readability and language. After using this tool, the authors thoroughly reviewed and modified the manuscript and took full responsibility for the content of the publication. The literature survey was done only by the authors, with no input from the AI tools.

## Authors’ Contributions

AZ: Conceptualization, Methodology, Software, Visualization, Writing – Original Draft, Writing – Review & Editing; MMZ: Conceptualization, Methodology, Writing – Review & Editing; AD: Investigation, Writing – Review & Editing; ASCz: Investigation, Writing – Review & Editing; PJ: Conceptualization, Supervision, Project administration, Funding Acquisition, Writing – Review & Editing.

## Additional Files

**Supplementary Table S1.** Description of sequencing datasets.

**Supplementary Table S2.** StringTie2 parameters tested on SIRVs.

**Supplementary Table S3.** Custom helper scripts used in the SmedAnno workflow

